# Fast and robust metagenomic sequence comparison through sparse chaining with skani

**DOI:** 10.1101/2023.01.18.524587

**Authors:** Jim Shaw, Yun William Yu

**Affiliations:** Department of Mathematics, University of Toronto; Computer and Mathematical Sciences, University of Toronto at Scarborough

## Abstract

Sequence comparison algorithms for metagenome-assembled genomes (MAGs) often have difficulties dealing with data that is high-volume or low-quality. We present *skani* (https://github.com/bluenote-1577/skani), a method for calculating average nucleotide identity (ANI) using sparse approximate alignments. skani is more accurate than FastANI for comparing incomplete, fragmented MAGs while also being > 20 times faster. For searching a database of > 65, 000 prokaryotic genomes, skani takes only seconds per query and 6 GB of memory. skani is a versatile tool that unlocks higher-resolution insights for larger, noisier metagenomic data sets.

---

High-throughput metagenomic sequencing is now generating terabytes of data [1], rendering many traditional workflows unusable. Consider the fundamental problem of computing sequence-to-sequence similarity between metagenome-assembled genomes (MAGs). Modern studies are generating hundreds of thousands of MAGs [2], and searching these MAGs against a database or computing all pairwise similarities takes billions of comparisons; this is infeasible with traditional alignment-based methods. Thus, large-scale sequence comparison for metagenomic data is now dominated by *sketching* methods [3,4]. Sketching methods summarize data sets into small collections of k-mers, known as *sketches*. These sketches can then be efficiently compared against one another.

Sketching methods such as Mash [3] or sourmash [4] approximate average nucleotide identity (ANI) through certain k-mer statistics (known as *indices*) built around estimating the overlap between the k-mers in two genomes, appropriately normalized. More precisely, these indices take the form 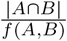 where *A* and *B* are the sets of k-mers in two genomes, and *f* (*A, B*) controls the normalization. The Jaccard index used in Mash [3] corresponds to *f* (*A, B*) = |*A ∪ B*|, and the containment index is *f* (*A, B*) = |*A*| [5]. The main index we will consider is the *max-containment index*, which corresponds to *f* (*A, B*) = min(|*A*|, |*B*|); minimizing the denominator maximizes the containment. While these indices themselves do not represent an ANI, they can be translated into an ANI through simple formulas [6,7].

All of the above k-mer statistics suffer from the following problem: as two MAGs become more incomplete, the overlap |*A ∩ B*| may decrease more than *f* (*A, B*) because a homologous k-mer is simply not present in the assembly, therefore making the index smaller. The decrease has nothing to do with the *genetic distance* between the genomes; it is simply an *artefact* of the assembly being incomplete. Even “medium-quality” MAGs typically only require > 50% completeness [2], so sketching methods can underestimate sequence identity; this has been shown to be problematic in practice [8]. On the other hand, alignment-based methods are able to pick out only the orthologous regions to estimate ANI from, so incompleteness is not an issue. Alignment also gives the *aligned fraction* (AF), the fraction of the genomes actually aligned to one another, which is a useful statistic that pure sketching methods do not estimate. There is thus a need for algorithms that are fast, like sketching methods, yet robust to noise due to assembly artefacts, like alignment methods.

These design constraints led to the earlier development of FastANI [9], a hybrid alignment and sketching method based around minimizer seeds [10]. However, FastANI is held back by the use of minimizer seeds, which are known to be biased when sparse [7]; thus, FastANI must use dense, slow seeding schemes. Fortunately, recent advances in the theory of k-mer seeding have led to the invention of more principled seeding schemes [6] that give theoretically unbiased ANI estimates.

We developed *skani*, a fast, robust tool for calculating ANI and aligned fraction. skani’s ANI method is robust against incomplete and fragmented MAGs, yet it is multiple orders of magnitudes faster than alignment-based methods and over an order of magnitude faster than even the state-of-the-art FastANI. skani uses a very sparse k-mer chaining [11] procedure to quickly find orthologous regions between two genomes. This allows for sequence identity estimation using k-mers on *only the shared regions* between two genomes (see Fig. 1a), avoiding the pitfalls of alignment-ignorant sketching methods. Like BLAST-based ANI methods, skani breaks genomes into non-overlapping fragments, estimates the ANI for each fragment, and then averages the ANI to output an ANI estimate. We then use a trained regression model to debias our ANI estimates (see Methods). This last step is optional, but enabled by default.

**Fig. 1:**
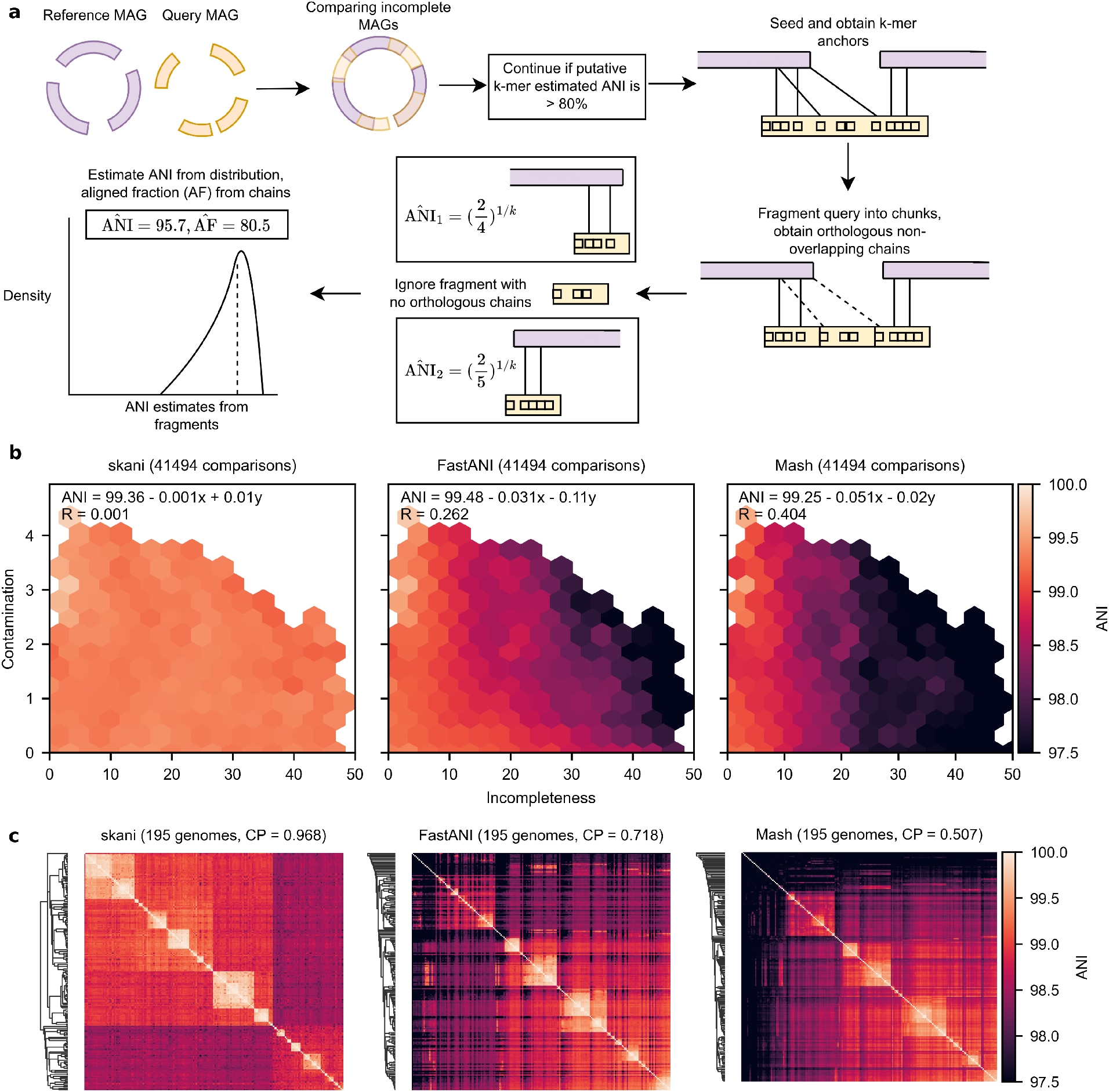
**a**. Algorithm overview of skani. **b**. ANI sensitivity to contamination and incompleteness. We took all pairs of MAGs with > 99% ANI according to ANIm from species-level bins generated by Pasolli et al. [2] with > 25 and < 50 genomes, leading to a diverse set of 41949 pairs of genomes. We re-evaluated the ANI of each method and performed ordinary least-squares regression with incompleteness and contamination (averaged between the pair and obtained by CheckM [12]) as covariates. Estimated parameters and *R*^2^ values are shown; only hexagons with > 20 data points are visible (see Extended Data Fig. 4 for density information). We also used N50 instead of incompleteness as a covariate in Extended Data Fig. 3. **c**. Average-linkage cluster heatmap for each method on bin number 2328 from Pasolli et al. [2] (classified as *Alistipes ihumii*) with 195 genomes. Cophenetic correlation (CP) of each method’s dendrogram (with ANIm’s distance matrix as a ground truth) is shown. skani’s high cophenetic correlation indicates that its dendrogram is concordant with ANIm’s dendrogram, which we show in Extended Data Fig. 1.

We first verified that existing ANI methods are indeed sensitive to incompleteness and fragmentation in a simulated test: in Extended Data Fig. 2, we simulated fragmentation and incompleteness for an *E. coli* genome to test ANI calculation robustness against assembly artefacts. Of course, true ANI values for these fragmented, incomplete, but *identical* genomes should be 100%. However, ANI estimates were systematically lower than 100%. Mash was the most affected: comparing two 50% complete genomes could lead to an average estimate of only 96%. FastANI also had issues on this simple test, as it was sensitive to fragmentation (i.e. low N50), which is why a minimum N50 of 10,000 is used in the original study [9], but that N50 requirement is not met in many real experiments [2,13]. The most robust method was ANIm [14], a MUMmer [15] based ANI method, so although it is too slow for many applications, we chose ANIm as a baseline for comparing MAGs.

Next, we showed on *real* MAGs that only skani and ANIm are robust to MAG quality for high-resolution ANI calculations. In Fig. 1b, we used MAGs and species-level bins generated by Pasolli et al [2], a large collection of medium-quality and high-quality short-read assembled MAGs. We ran all ANI methods on an arbitrary set of 41,494 pairs of genomes where the ANIm predicted ANI was > 99%. As contamination and incompleteness increases, FastANI and the sketching-only methods have a visible trend toward lower ANI values. However, ANIm’s ANI calculations in Extended Data Fig. 1 show no such trend, suggesting that these decreases in ANI are spurious. Extended Data Fig. 3 shows the primary cause for FastANI’s decrease in ANI is low N50, agreeing with our simulations. We quantified this effect by regressing over incompleteness and contamination as covariates. The low *R*^2^ values of ANIm (0.005) and skani (0.001) confirm the qualitative observation that skani is robust towards noise, whereas all other methods except ANIm have *R*^2^ > 0.2.

Because ANI underestimations due to MAG quality are *systematically biased*, such ANI estimates can strongly impact phylogenetic observations. We show in Fig. 1c that dendrograms (obtained by average-linkage clustering) for a species-level bin differ greatly between ANI methods. skani’s dendrogram qualitatively resembles ANIm’s dendrogram (Extended Data Fig. 1) more closely than the other methods, yet it is > 500 times faster than ANIm and > 50 times faster than FastANI for computing the distance matrix (Extended Data Fig. 5).

To quantify the goodness of these dendrograms, we use cophenetic correlation [16]. Given any hierarchical clustering dendrogram and any distance matrix, cophenetic correlation gives a value from -1 to 1, with higher values indicating stronger concordance between the dendrogram and matrix. We evaluate each method’s dendrogram, obtained by average-linkage clustering from its *own distance matrix*, against *ANIm’s distance matrix*. As a baseline, we take the correlation between ANIm’s dendrogram against its own distance matrix.

We verify in Extended Data Fig. 6 using cophenetic correlation that skani’s clusterings are indeed more concordant with ANIm than other methods. skani’s clusterings agree with ANIm’s baseline well (−0.039 mean deviation) on a set of 238 species-level bins from Pasolli et al, whereas FastANI and Mash have a much larger mean deviation (−0.228 and -0.424 respectively). Interestingly, using the max-containment index with sourmash led to a better mean deviation (−0.148), but this is an option that has to be specified, and it is still several times worse than skani. Notably, the ordering of the methods with respect to the mean cophenetic correlation is skani > sourmash max-contain > FastANI > mash; this exactly agrees with the ordering using *R*^2^ values from the contamination-incompleteness plots in Fig. 1b and Extended Data Fig. 1 – implying that MAG assembly artefacts are indeed to blame for the clustering discordance.

In Extended Data Fig. 11 - 14, we explore skani on three additional data sets: ocean eukaryotic MAGs, [17,18], ocean archaea MAGs [13], and soil prokaryotic MAGs [19], which in addition to the Pasolli et al data set gives four data sets in total. We describe these data sets in Supplementary Table 2 as well.

Extended Data Fig. 11 shows that skani’s results generalize to a diverse set of genomes, including *eukaryotic* MAGs with an N50 of 17.6 Mbp (and 32 MAGs of size > 100 Mbp). When comparing each method’s deviation from ANIm and considering the 1 to 99 percentile deviations, skani has the smallest 1 to 99 percentile interval lengths for all data sets, indicating robustness. We also show in Extended Data Fig. 12 and 13 that skani has a better linear aligned fraction correlation with ANIm than FastANI on every data set. The accuracy can be improved further by controlling subsampling rate of the k-mers (parameter *c* in Methods), in which case the Pearson R is > 0.996 on three of the four data sets, showing that skani’s AF is almost in perfect concordance with MUMmer.

An important task is *classifying* MAGs (or isolate genomes) by searching against a database of reference genomes. Such databases represent a diverse collection of genomes where only a fraction of the genomes are similar to the query. Therefore, sensitively searching against each reference is unnecessary. To enable efficient database search, we augmented skani with a near constant-time ANI filter against distant genomes.

skani’s fast ANI filter first computes the max-containment index for a very sparse set of *marker* FracMin-Hash k-mers to approximate ANI [6]. We perform sensitive calculations only if the estimate is higher than a threshold. Importantly, the sparsity of the marker k-mers (roughly 1 out of 1000 k-mers chosen by default) means we can insert all marker k-mers into a hash table as keys with values corresponding to the genome containing the marker (i.e. an *inverted index*). This makes calculating the max-containment for a query by hashing all of its marker k-mers take essentially constant time when the database is diverse. skani can choose to only load a genome’s seeds fully into memory if the genome passes the filter, allowing for memory-efficient search. By default, we enable the hash table filter if there are > 50 query genomes, otherwise, a simple iterative filter is slightly faster than building the table.

We show in Fig 2 that skani can query an E.coli genome against the GTDB database [20] (> 65, 000 genomes) in seconds using < 6GB of RAM, which is comparable in speed to Mash. skani is much faster than FastANI for querying (> 20 times on the *E. coli* data set) and is > 2.5 times faster than Mash for indexing. Furthermore, skani can do all-to-all comparisons on a set of 4233 RefSeq complete representative bacterial genomes as quickly as Mash due to the fast filtering of unrelated genomes. FastANI takes much longer (Extended Data Fig. 9), but the comparison is unfair to FastANI due to FastANI allowing lower ANI genome comparisons than skani and thus doing more computations. In Extended Data Fig. 7 we show an additional three experiments, and all data sets are described in Supplementary Table 2. A drawback of skani is that the index file sizes are larger than pure sketching methods (Supplementary Table 3), but disk space is usually less of an issue than RAM.

**Fig. 2:**
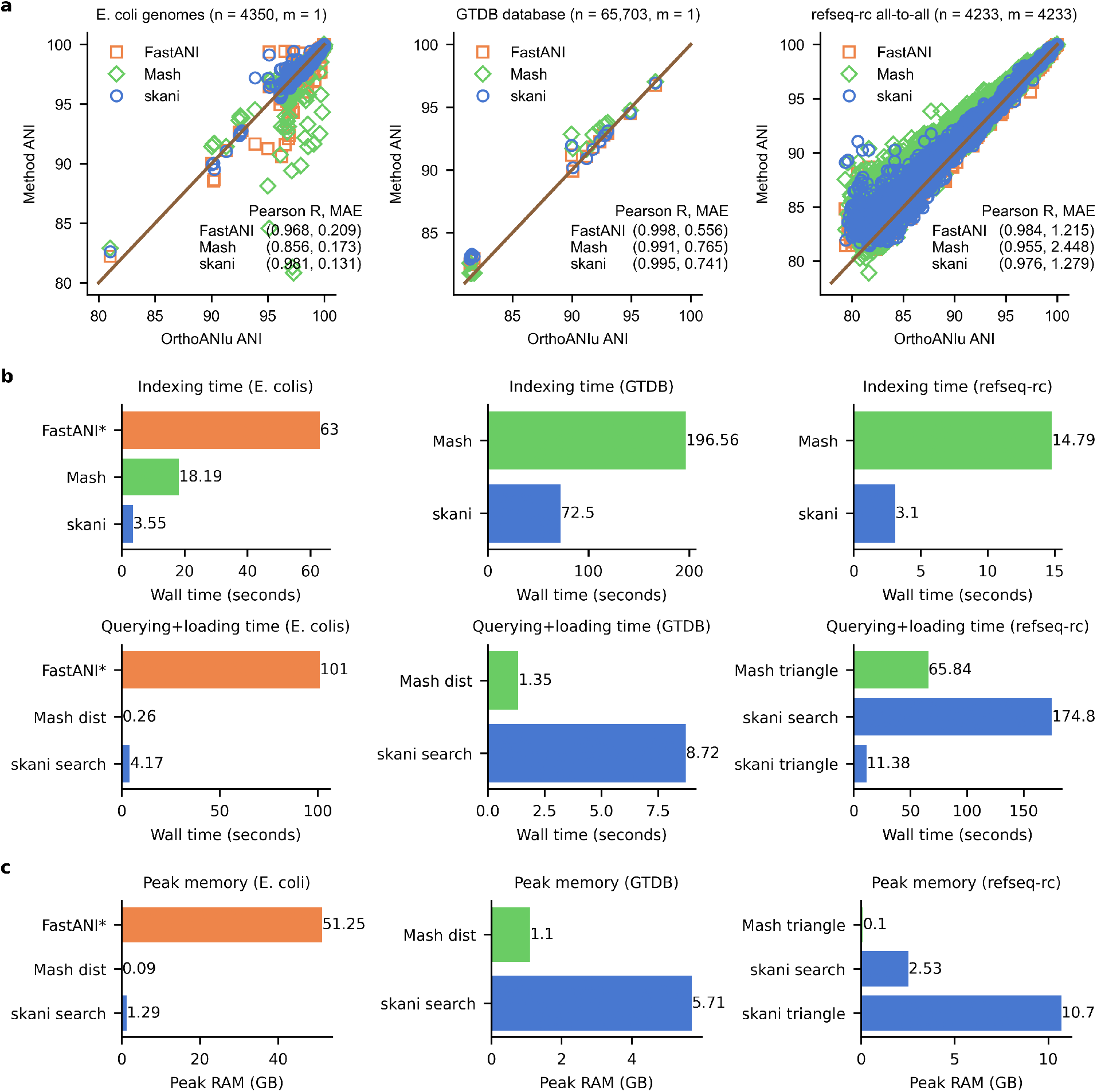
**a**. ANI benchmarking with *n* reference genomes and *m* query genomes. From left to right: (1) querying a single *E. coli* K12 genome against a collection of *E. coli* genomes, (2) querying an *E. coli* genome against the GTDB database, and (3) all-to-all comparisons on the refseq-cr (complete and representative bacterial genomes) database. BLAST-based ANIu [21] is used as a baseline. We only analyzed data points for which all methods had a predicted value. Pearson R value and mean absolute error (MAE) are shown for each data set. Data set descriptions can be found in Supplementary Table 2. **b**. Indexing and querying wall time for each data set (50 threads). See Extended Data Fig. 8 for CPU times instead. Subcommands used are shown for each method when applicable. FastANI indexing and query times were estimated from the output. skani search and skani triangle are different subcommands that give the same results, but skani search only loads genomes into RAM as needed and discards after usage. FastANI times are not shown for the GTDB and refseq-cr data sets for fairness to FastANI due to FastANI’s ability to output a slightly larger range of ANI values (approximately > 75% for FastANI vs > 80% for skani). **c**. Peak memory usage for each method and subcommand. Sketching took negligible memory for skani and Mash.

In Fig. 2 a and Extended Data Fig. 7, skani’s accuracy is usually better than Mash but slightly worse than FastANI for reference-quality genomes, with the difference between skani and FastANI in mean absolute error (MAE) never more than 0.20 ANI. skani is better on the *E. coli* data set, which includes many fragmented, possibly incomplete genomes that give rise to many Mash and FastANI outliers. The results for FastANI improved (Pearson R from 0.974 to 0.994) on the *E. coli* data set if we removed genomes with N50 <10,000, which is the same as the originally reported FastANI results [9]. Thus skani gives only slightly less accurate values than FastANI on reference-quality genomes with the assurance of robustness for low-quality assemblies.

We have shown that skani improves on the state-of-the-art for metagenomic sequence comparison. skani is almost as fast as sketching-based methods for ANI database search, yet it gives a more robust phylogenetic signal when comparing noisy MAGs. Given the overwhelming amount of data generated by modern metagenomic studies, we believe skani’s ability to analyze an order of magnitude more data *while simultaneously* giving a stronger signal will allow examination of vast metagenomic sequences at a higher resolution, unlocking new types of analysis not possible before.

## Methods

### Sequence identity estimation

Formally, let *G* be a string of nucleotides and *G*^*′*^ be a mutated version of *G* where every letter is independently changed to a different letter with probability *θ*. We will define the true ANI to be equal to 1 *− θ* under our model. Under the usual assumption of no repetitive k-mers [22], it is easy to estimate *θ* from k-mer matching statistics [6,22,3] between *G* and *G*^*′*^. We proved in a previous work that for random, mutating strings, the expected value of such spurious matches for a string of length *n* is 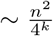 (Theorem 2 in Shaw and Yu [23]), so this is not a bad assumption in practice when *k* is reasonably large and for simpler, non-eukaryotic genomes. However, when dealing with MAGs, we don’t have *G* and *G*^*′*^ but instead fragmented, contaminated, and incomplete versions of *G* and *G*^*′*^. The models used in these sketching methods give biased estimates for ANI that are too small because missing k-mer matches may be due to mutations, incompleteness, or contamination instead of *only* due mutations. Thus to accurately use k-mer statistics, we first find orthologous regions by approximate alignment, and then use k-mer statistics.

### Algorithm outline

The main idea behind skani is to find an approximate set of orthologous alignments between two genomes by obtaining a set of non-overlapping k-mer chains [11] (the *chains* don’t overlap; the *k-mers* may overlap within the chain). We can then estimate ANI from the statistics of the k-mers in the chains, avoiding costly base-level alignment. The main algorithmic steps are listed below.

1. We use a very sparse set of *marker ℓ*-mers to estimate max-containment index and obtain a putative ANI. We filter out pairs of genomes with putative ANI < 80%.
2. We select the genome with the larger score, defined as total sequence length times mean contig length, to be the reference and the other to be the query. We then fragment the query into 20kb non-overlapping chunks.
3. 3.We extract 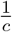 fraction of k-mers for both genomes for some *c* (*c* = 125 by default) as *seeds* to be used for chaining.
4. For each chunk on the query, we chain the seeds using a standard banded, heuristic chaining method against the reference.
5. We greedily extract minimally-overlapping chains between the query and the reference and output aligned fraction.
6. We estimate the ANI for each chunk, and output the mean ANI over all chunks.

### Sketching by FracMinHash

Instead of using the set of all k-mers in a genome, we use a compressed representation by *sketching*, i.e. selecting only a subset of all k-mers. To obtain such a set of k-mers, we use the FracMinHash method [24]: given a hash function *h* which maps k-mers to [0, *M*], we select the k-mer *x* as a seed if *h*(*x*) *< M/γ*, where *γ* controls the fraction of selected k-mers. Assuming a uniform hash function, the expected fraction of selected k-mers is 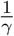.

While FastANI uses minimizer [10] k-mers to estimate Jaccard index and then ANI, recently it was shown that Jaccard estimates (and thus ANI estimates) from minimizer k-mers are biased [7] and depend crucially on the window size *w*. While this bias is not too bad when the *w* is small (FastANI uses relatively small *w* = 24), it scales as *w* increases. This means that a minimizer ANI estimator can not use very sparse seeds, since the fraction of selected seeds is 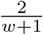[25].

### Max-containment putative ANI screening with marker *ℓ*-mers

skani is not optimized for comparing distant genomes, so we can filter out comparisons against distant genomes using a very sparse set of FracMinHash *ℓ*-mers, which we call *markers* (terminology inspired from Shafin et al. [26]). We use these markers to estimate the *max-containment index* [5]; the same method is implemented in sourmash [6], although sourmash does not use max-containment by default. Let a set of markers obtained from FracMinHash from the genome *G*_1_ (with *γ* = *c*_*m*_ and *ℓ*-mers) be denoted as *A*, and denote *B* as the analogous set from the genome *G*_2_. We then calculate an ANI estimate between two genomes *G*_1_, *G*_2_ as

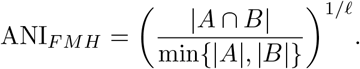

The term on the right inside the exponent is the max-containment index. FracMinHash has only negligible bias in calculating the max-containment index [6], and can be used to obtain an estimate of ANI no matter the density (i.e. the fraction of selected k-mers). We let *ℓ* = 21 for our marker *ℓ*-mers and *c*_*m*_ = 1000 by default. We found skani’s ANI algorithm was most accurate when ANI ≥∼ 82%, so we only compare genomes with ANI_*F MH*_ > 80% as a conservative underestimate.

We can store all markers contained in the reference genomes as keys in a hash table with the associated values being the set of genomes containing the key. Given a query genome, we can then obtain the intersection of all markers for the query against the references in running time only dependent on the number of similar genomes in the references [27]. From this, we can obtain ANI_*F MH*_, making our filtering step truly sub-linear as long as our reference and query genomes are diverse. When performing all-to-all pairwise comparisons, skani uses this strategy by default. However, when querying a small number of genomes against a database, skani iteratively checks all pairs of genomes by default because building the hash table takes a non-trivial amount of time.

### Obtaining sparse seeds for chaining

The marker *ℓ*-mers described above are only used for filtering, but not for actually estimating the ANI. To actually estimate the ANI, we use a hybrid approach via local mapping (without base-level alignment) *and* containment index estimation. The first step in our approach is to obtain a *different* set of FracMinHash k-mers, this time taking *γ*, the sampling rate of k-mers, to be *γ* = *c* where *c* < *c*_*m*_, and *k* < ℓ. This gives a denser, more sensitive set of k-mers to be used as *seeds*. These new k-mers are called seeds instead of markers because we actually use them as seeds for k-mer matching and alignment. We note that while we could have used other “context-independent” k-mer seeding methods that are more “conserved” than FracMinHash [28], we found that FracMinHash works well enough for relatively sparse seeds when *c* ≫ *k*. By default, *k* = 15 and *c* = 125. We note that the small value of *k* = 15 used by default leads to too many repetitive anchors on larger genomes, so we mask the top seeds that occur more than 2500/*c* times by default. See the section *Chaining score function and algorithm* below for a justification of the choice of 2500/*c*.

### Chaining sparse k-mer seeds

After selecting one genome as a reference and the other as the query, we fragment the query into 20kb non-overlapping chunks. For each chunk, we find a set *anchors*, which are exact k-mer matches between *g* and *g*^*′*^. Each anchor can be described by a tuple (*x, y*), indicating the starting position of the matching k-mers on *g* and *g*^*′*^ respectively. We collect the anchors into a strictly increasing subsequence called a *chain* [11] based on the ordering (*x*_1_, *y*_1_) *≺* (*x*_2_, *y*_2_) if *x*_1_ < *x*_2_ and *y*_1_ < *y*_2_.

We sort the anchors (*x*_1_, *y*_1_), … (*x*_*N*_, *y*_*N*_) in lexicographic order and let *S*(*i, j*) be the score of chaining the *i*th anchor to the *j*th anchor. We wish to find a strictly increasing subsequence (based on our previously defined ordering ≺) of anchors (*i*_1_, *i*_2_, …) maximizing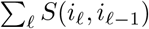. The optimal such subsequence can be calculated in *O*(*N* ^2^) time where *N* is the number of anchors by the following dynamic programming: letting *f* (*i*) be the optimal score of the chain up to the *i*th anchor, let

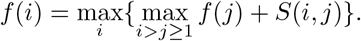

After calculating *f* (*i*) for each anchor *i*, we can obtain a set of optimal chains. We describe this in detail below.

### Chaining score function and algorithm

Letting (*x*_*i*_, *y*_*i*_) and (*x*_*j*_, *y*_*j*_) be the *i*th and *j*th anchors, we define our chaining cost to simply be *S*(*i, j*) = 20 − |(*y*_*j*_ − *y*_*i*_) − (*x*_*j*_ − *x*_*i*_)|. Notably, we allow and do not penalize overlapping k-mers because this would bias the ANI calculation, as we want the chain to include all k-mers that arise from sequence homology. We also don’t need to use more sophisticated scoring functions because we do not actually need to worry about base-level alignments or finding the longest chain. Furthermore, we use a banded dynamic programming method where instead of iterating over all *j < i*, we iterate only up to *i* − *A* < *j* < *i* or stop if |*x*_*i*_ − *x*_*j*_| > *B* for some constants *A* and *B*. Thus the worse case time is *O*(*AN*) instead of *O*(*N* ^2^). By default, we let *A* = *B/c*, where *B* = 2500 and *c* is the seed subsampling rate.

### Obtaining homologous chains by backtracking

For each chunk, after computing all optimal scores *f* (*i*) over all anchors using banded dynamic programming, we obtain optimal chains using the standard method of backtracking. That is, we store an array of pointers corresponding to anchors, where the pointer points to the optimal predecessor determined by the dynamic programming. For any anchor, we could then trace through this array to obtain the best chain corresponding to each anchor.

We wish to obtain a set of chains that don’t share any anchors for each chunk. To do this, we partition the anchors into disjoint sets using a union-find data structure, taking unions of two anchor representatives whenever one anchor is an optimal predecessor of another. We then find the best *f* (*i*) within each disjoint set and backtrack to optimal the optimal chain within each disjoint set. We take set of all such chains over *all chunks* and call these chains *homologous chains*.

### Obtaining orthologous chains from homologous chains

It is possible that a single region chains to multiple paralogs, so the homologous chains obtained above may not be orthologous. To obtain orthologous chains, we use a greedy minimally-overlapping chain-finding procedure. We first sort every homologous chain (over *all chunks*) by its score and examine the best chain, only examining chains with ≥ 3 anchors. If the overlap between this current best chain and any other already selected chains is less than 50% of the current chain’s length, we select this best homologous chain as an orthologous chain and then examine the next best homologous chain. We repeat this procedure until all remaining homologous chains overlap an orthologous chain by more than 50% and return all orthologous chains.

This procedure is similar in spirit to the reciprocal mapping method used in other ANI methods [21,9] to capture orthology, but it avoids performing alignment twice, making running time twice as fast. Other more sophisticated methods [29] exist for finding sets of orthologous alignments but we found this heuristic to be good enough for our method.

### Estimating ANI from chains

For a given chunk, let *α* be the number of anchors in the non-overlapping orthologous chains on that chunk, and *M* be the number of seeds in the chunk. Then we estimate an ANI for each chunk by

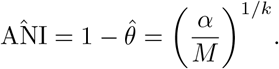

This comes from modelling each k-mer as being an exact match and thus an anchor with probability (1 − *θ*)^*k*^ under a simple, independent mutation model, so the expected number of anchors is (1−*θ*)^*k*^ times the number of k-mers [6].

However, the above formula fails when only part of the chunk is homologous to the reference, which may happen due to incompleteness of MAGs or structural variation. This will underestimate the ANI, since not all seeds in the chunk arise from sequence homology. Let *M*_*LR*_ be the number of seeds in the chunk contained only between the leftmost and rightmost anchor over all non-overlapping chains, and consider

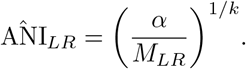

If 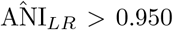 and the number of bases covered by in between these flanking anchors is > 4 *∗ c*, then we use 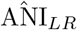 as the estimate instead. This heuristic comes from the observation that when the ANI is large enough, our sparse chains are relatively accurate, so we can truncate the chunk with good accuracy. However, if ANI is small, it is more possible that this heuristic incorrectly excludes seeds that arise from sequence homology, so we avoid applying the heuristic to distant pairs of genomes.

### Estimating final ANI and AF

To obtain our final ANI estimates, we take a weighted mean over all chunks with weights given by *M* (or *M*_*LR*_ if the heuristic is applied instead), the number of seeds in the chunk. We also report the reference and query alignment fraction as the sum of the bases covered by all chains divided by the respective genome size. The number of bases in a chain is the last anchor position minus the first anchor position plus 2 *∗ c*, where the 2 *∗ c* term is to account for k-mers near the edge of the chain missed by subsampling. By default, we only output a final ANI value if the alignment fraction is greater than 15%.

We optionally output a 90% confidence interval for skani by performing a percentile bootstrap on the ANI distribution over the non-overlapping chunks. We use 100 iterations by default, and only proceed if there are > 10 chunks. We found that the bootstrap gave reasonable ANI confidence intervals over *skani’s inherent randomness*, i.e. stochasticity due to k-mer sampling, but we stress that it does not account for any systematic biases of skani’s ANI estimator relative to other methods.

### Non-parametric regression for ANI debiasing

The chaining procedure can overestimate ANI because the chains may exclude homologous but mutated k-mers near the edges of the chain due to the local mapping procedure. To handle this, we introduce an optional post-processing regression step to debias skani. We show in Extended Data Fig. 10 that this debiasing procedure corrects the ANI to be closer to a MUMmer-based ground truth.

We trained a gradient boosted regression tree with absolute deviation loss where the target variable is ANIm’s ANI calculation, and the features consist of the following: skani’s ANI, standard deviation of skani’s putative ANI distribution, and the 90th percentile contig lengths in the reference and query, and the average length of the k-mer chains, giving 5 features in total. We trained skani on a large, diverse set of 52,515 MAGs from Nayfach et al [30]. These assembled MAGs are disjoint from the other sets of MAGs used in this study. We computed all-to-all pairwise ANI values with > 90% ANI according to an untrained version skani, and ran ANIm on the resulting 1,004,213 pairs of genomes. To tune hyper parameters such as tree depth, number of trees, and learning rate, we chose a set of human gut MAGs [31] which is disjoint from the training set and the data sets used in our results, and then optimized our parameters over this new data set.

We only debias comparisons with putative ANI > 90% and > 150, 000 aligned bases. We enable the debiasing procedure by default when the parameter *c* is > 70. We noticed that for smaller *c*, bias is less of an issue. We trained two models, one for *c* = 125 (default) and another for *c* = 200. If the user selects a specific value of *c*, the model with corresponding *c* closer to the selected value is used. In particular, our results with *c* = 125 have debiasing enabled, but our results with *c* = 30 do not.

### Benchmarking details

All runtimes were benchmarked on a Intel(R) Xeon(R) CPU @ 3.10GHz machine with 64 cores and 240 GB of RAM as a Google Cloud instance with a persistent SSD disk. Unless otherwise specified, all programs were run using 50 threads. Exact commands are shown in Supplementary Table 1.

## Data availability

All data sets are specified in Supplementary Table 2.

## Code availability

skani is available at https://github.com/bluenote-1577/skani. Scripts and notebooks for reproducing all figures is available at https://github.com/bluenote-1577/skani-test.

## Acknowledgements

J.S was supported by an NSERC CGS-D scholarship. Work supported by Natural Sciences and Engineering Research Council of Canada (NSERC) grant RGPIN-2022-03074.

## Author contributions

J.S. conceived the project, designed the algorithms, and implemented skani. Y.W.Y supervised and contributed to the development of the methods. Both authors wrote and edited the manuscript.

---

**Supplementary Table 1:**
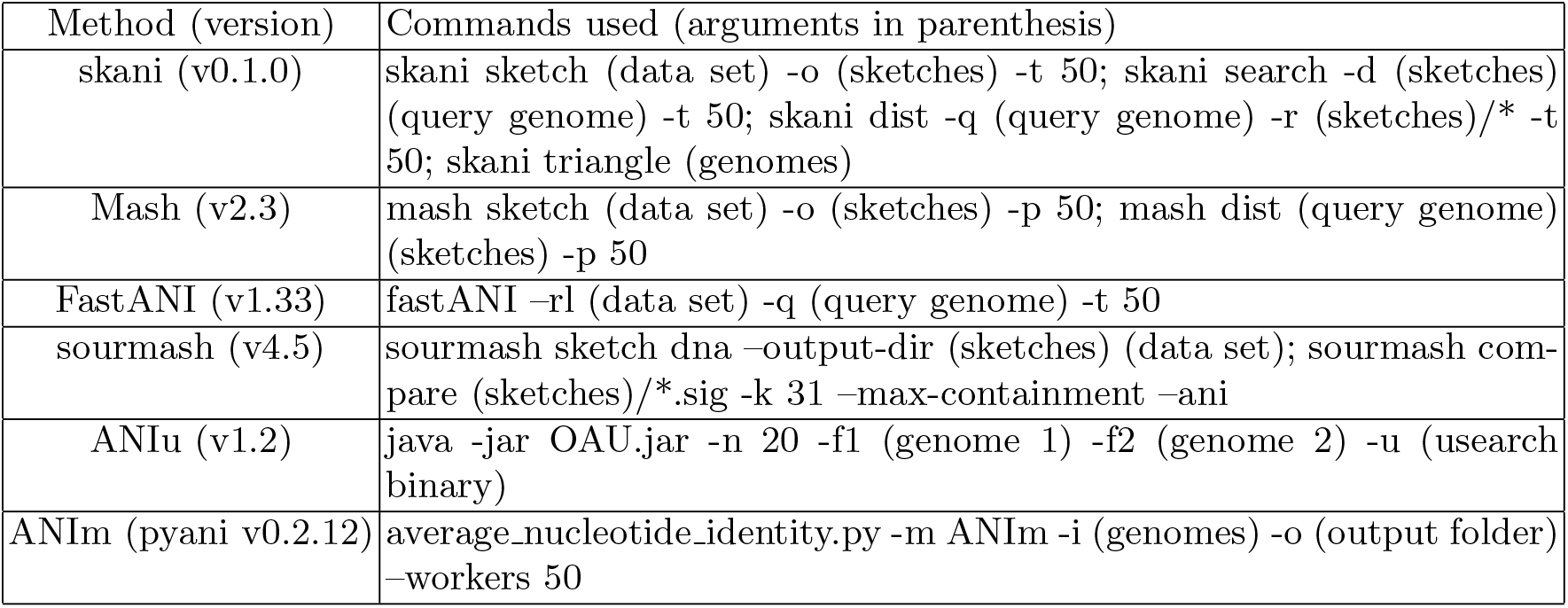
Method commands and parameters used for benchmarking.

**Extended Data Fig. 1:**
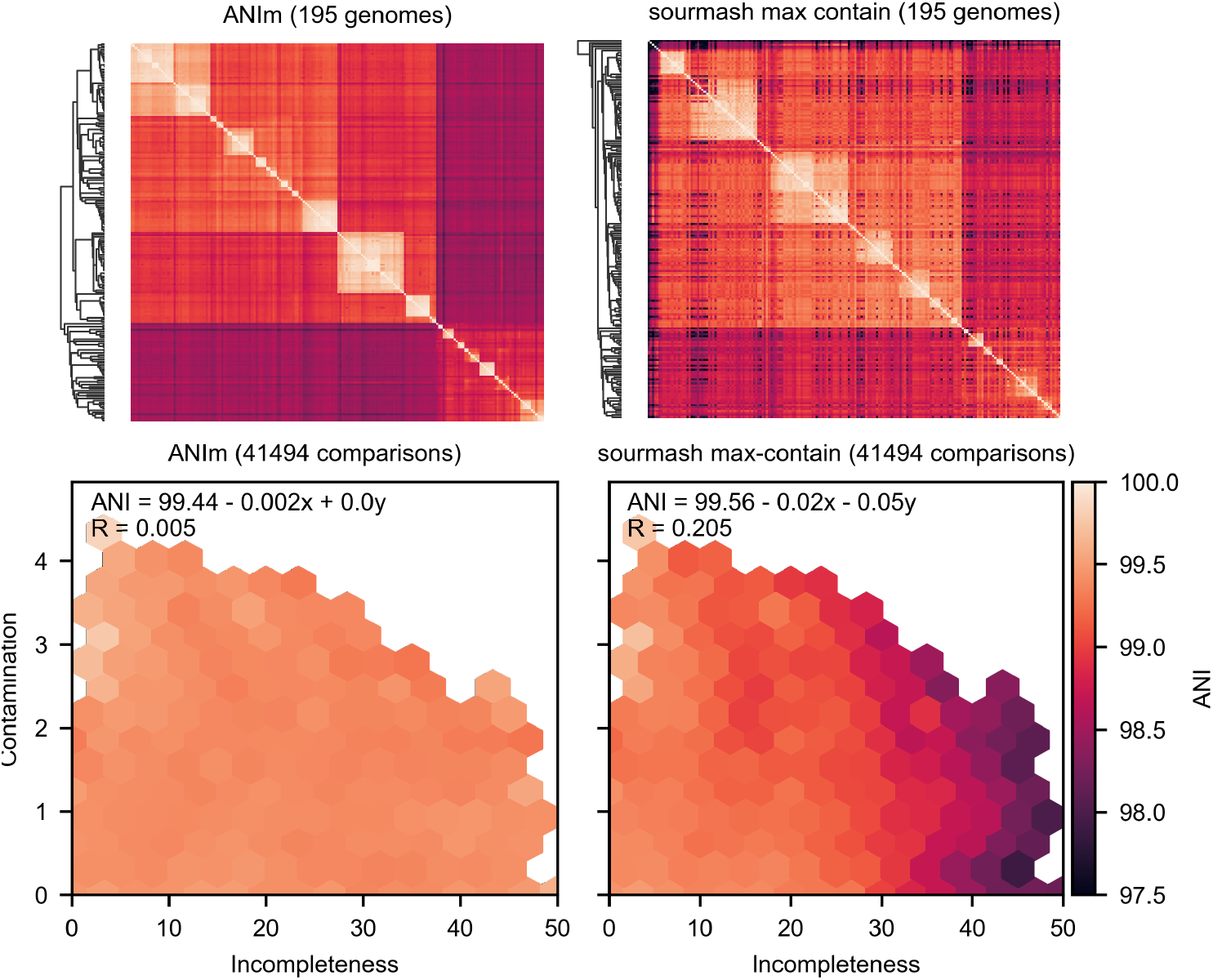
ANIm and sourmash max-containment clustering heatmaps and ANI versus incompleteness and contamination heatmaps corresponding to the experiments shown in Fig 1. sourmash max-containment estimates ANI using the max-containment index, corresponding to the --max-containment option in sourmash.

**Supplementary Table 2:** Extended description of data sets used in Fig. 1 and three additional data sets (Parks MAGs, *B. anthracis* genomes, and *B. fragilis* genomes) with additional plots shown in Extended Data Fig. 7. The refseq-cr and *B. fragilis* data sets were generated using ncbi-genome-download from https://github.com/kblin/ncbi-genome-download.

| Data set name | Description | Query genome(s) | Availability |
| --- | --- | --- | --- |
| Pasolli et al 25-50 | All MAGs in Pasolli et al [2] in species-level bins between 25-50 genomes (8611 genomes) | all-to-all comparisons within each bin | http://segatalab.cibio.unitn.it/data/Pasolli_et_al.html |
| Ocean archaea MAGs | 4435 archaea MAGs in non-representative OceanDNA MAGs from Nishimura and Yoshizawa [13] with > 90% ANI (according to skani) | all-to-all | https://doi.org/10.6084/m9.figshare.c.5564844.v1 |
| Soil MAGs | 1859 soil prokaryotic MAGs from soil genomes collected in Olm et al [19] with > 90% ANI (according to skani) | all-to-all | https://figshare.com/collections/Genomes_for_consistent_metagenome-derived_metrics_verify_and_define_bacterial_species_boundaries/4508162/1 |
| Ocean eukaryotic MAGs | 982 eukaryotic MAGs from Alexander et al [17] and 713 eukaryotic MAGs from Delmont et al [18] with > 90% ANI (according to skani) | all-to-all within each data set | https://osf.io/gm564/ and https://www.genoscope.cns.fr/tara/localdata/data/SMAGs-v1/SMAGs_contigs_individual.fna.tar.gz |
| Nayfach MAGs | 52,515 MAGs from Nayfach et al [30] | all-to-all | https://genome.jgi.doe.gov/GEMs |
| *E. coli* genomes | 4350 *E. coli* genomes. Data set D3 from Jain et al. [9]. | *E. coli* K12 (NC_007779) | Data set D3 from http://enve-omics.ce.gatech.edu/data/fastani. |
| *B. anthracis* genomes | 571 *B. anthracis* genomes. Data set D2 from Jain et al. [9]. | Bacillus anthracis 52-G (NZ_CM002395.1) | Data set D2 from http://enve-omics.ce.gatech.edu/data/fastani. |
| refseq-cr (complete + representative) | 4233 complete/chromosome level representative assemblies from refseq. Downloaded Sept. 16, 2022 | all-to-all | ncbi-genome-download --assembly-levels complete,chromosome --refseq-categories representative --formats fasta bacteria,viral,archaea,fungi |
| GTDB | 65703 genomes from the GTDB database [20], version 207. | *E. coli* K12 (NC_007779) | Download at https://gtdb.ecogenomic.org/ |
| Parks MAGs | 7901 MAGs from Parks et al. [32]. | *Pseudomonas stutzeri* (GCA_002292085.1) MAG (also from Parks et al. [32]) | Data set D5 from http://enve-omics.ce.gatech.edu/data/fastani. |
| *B. fragilis* genomes | 318 *B. fragilis* genomes from refseq. | *Bacteroides fragilis* YCH46 (NC_006347.1) | ncbi-genome-download --genera "Bacteroides fragilis" bacteria --formats fasta |

**Supplementary Table 3:** Size of stored index (i.e. sketches) on disk for three data sets.

| Data set | Size of index on disk |
| --- | --- |
| *E. coli* genomes (4350 genomes) | 4.1 GB |
| GTDB (65,703 genomes) | 40 GB |
| refseq-cr (4233 genomes) | 3.0 GB |

**Extended Data Fig. 2:**
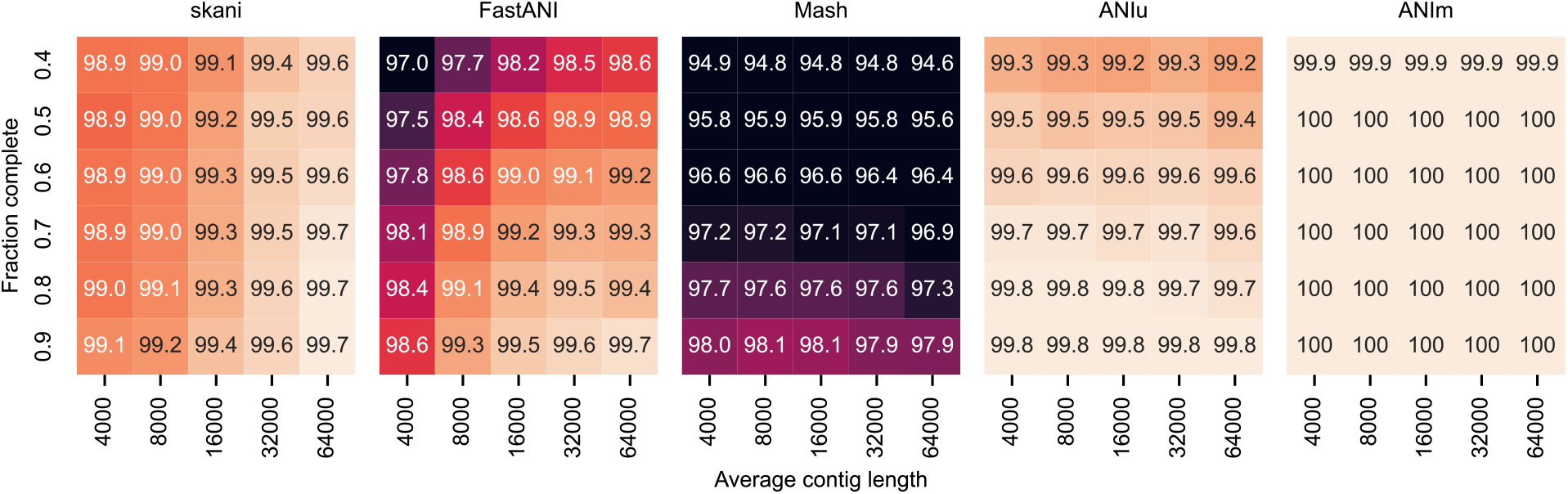
ANI benchmark under simulated fragmentation and incompleteness. We fragmented an *E. coli* genome to obtain non-overlapping contigs with lengths distributed according to an exponential distribution (mean length on the x-axis) and then retained each contig with some probability (probability on the y-axis). These simulated MAGs (20 for each pair length and probability parameters) were compared to each other and the average is shown.

**Extended Data Fig. 3:**
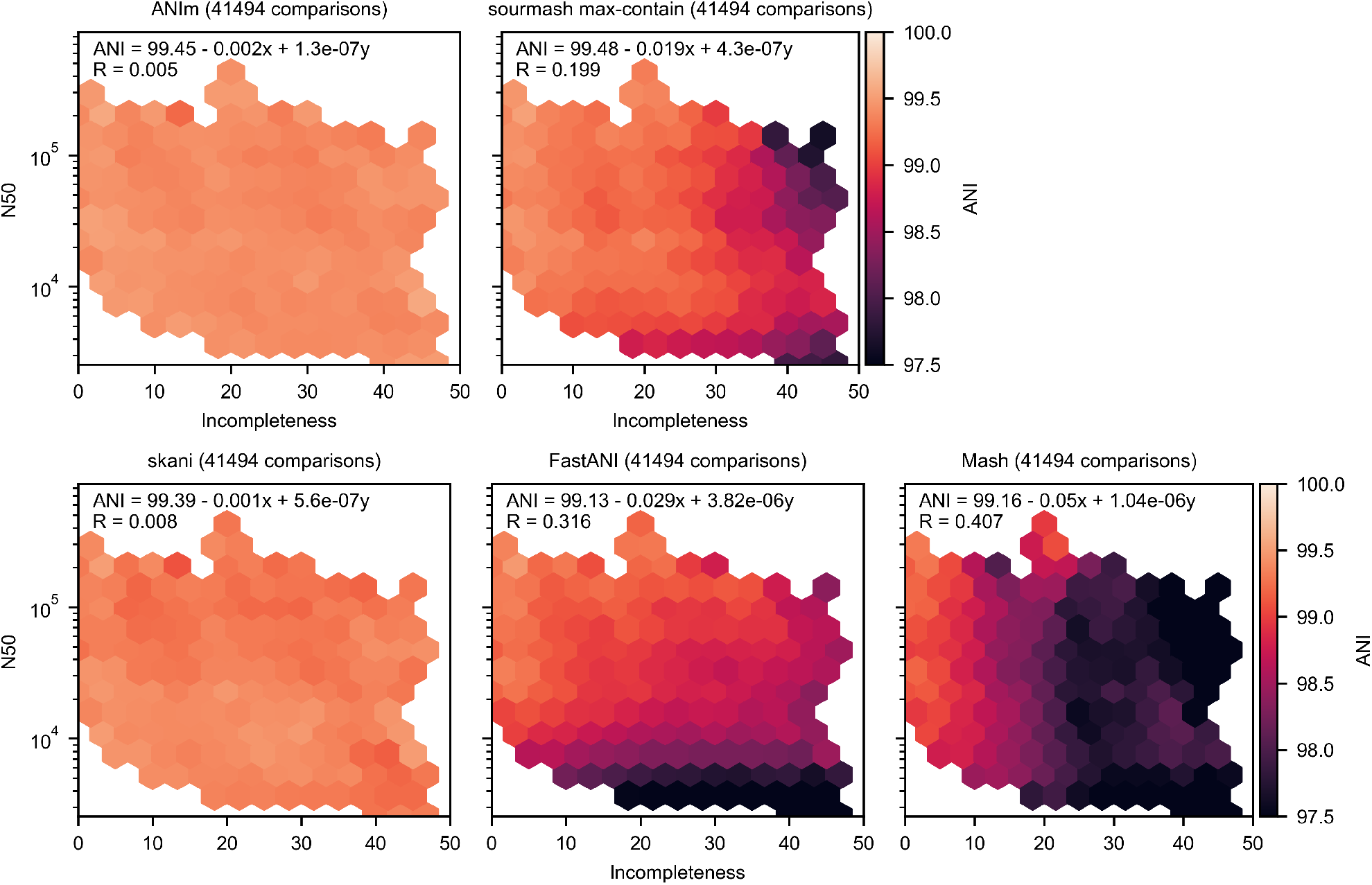
ANI sensitivity to fragmentation and incompleteness. The same experiment was run as in Fig. 1 b on Pasolli et al 25-50, except we used N50 as a covariate instead of contamination.

**Extended Data Fig. 4:**
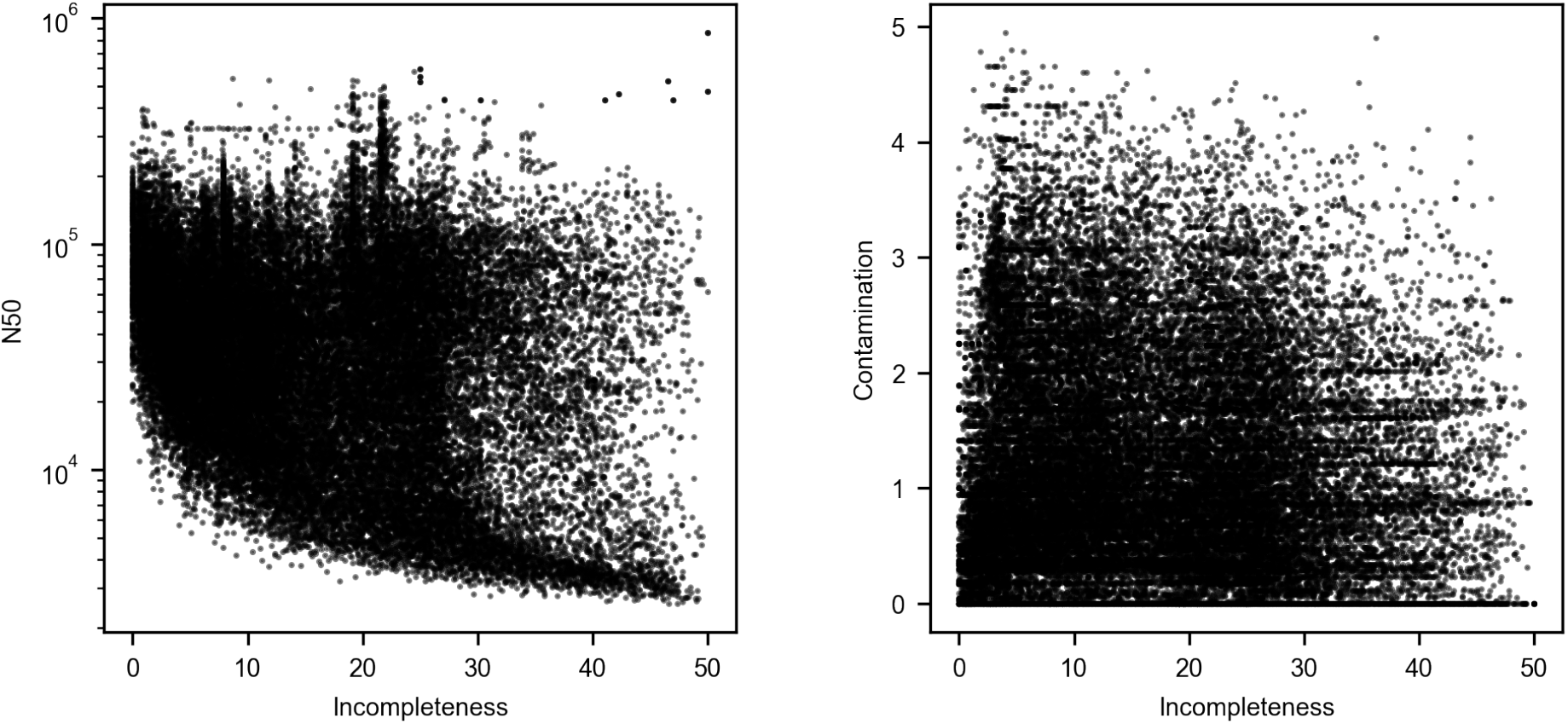
Scatter plot of the data points, representing pairs of genomes, in Fig 1 b and Extended Data Fig. 3.

**Extended Data Fig. 5:**
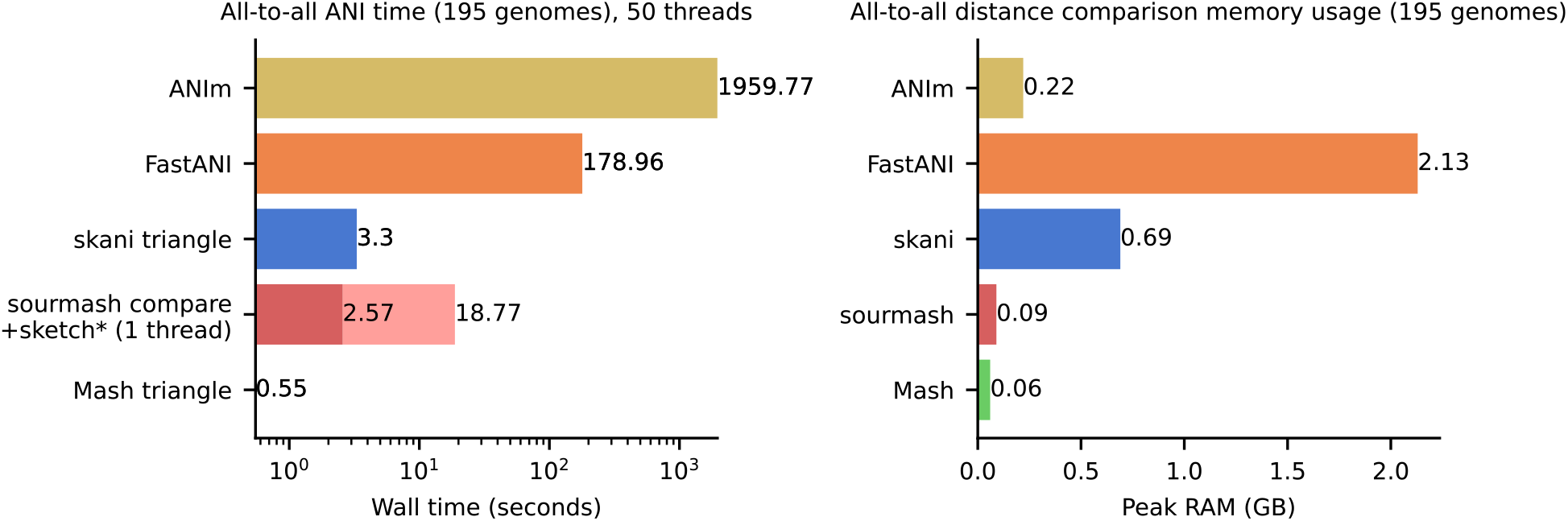
Running times and memory usage for all-to-all comparisons on the 195 genomes in Fig. 1 c with 50 threads. sourmash’s latest release does not have multi-threading support so sourmash was run with one thread. Out of sourmash’s 18.77 seconds runtime, 2.57 was used for ANI computation and the rest was for sketching.

**Extended Data Fig. 6:**
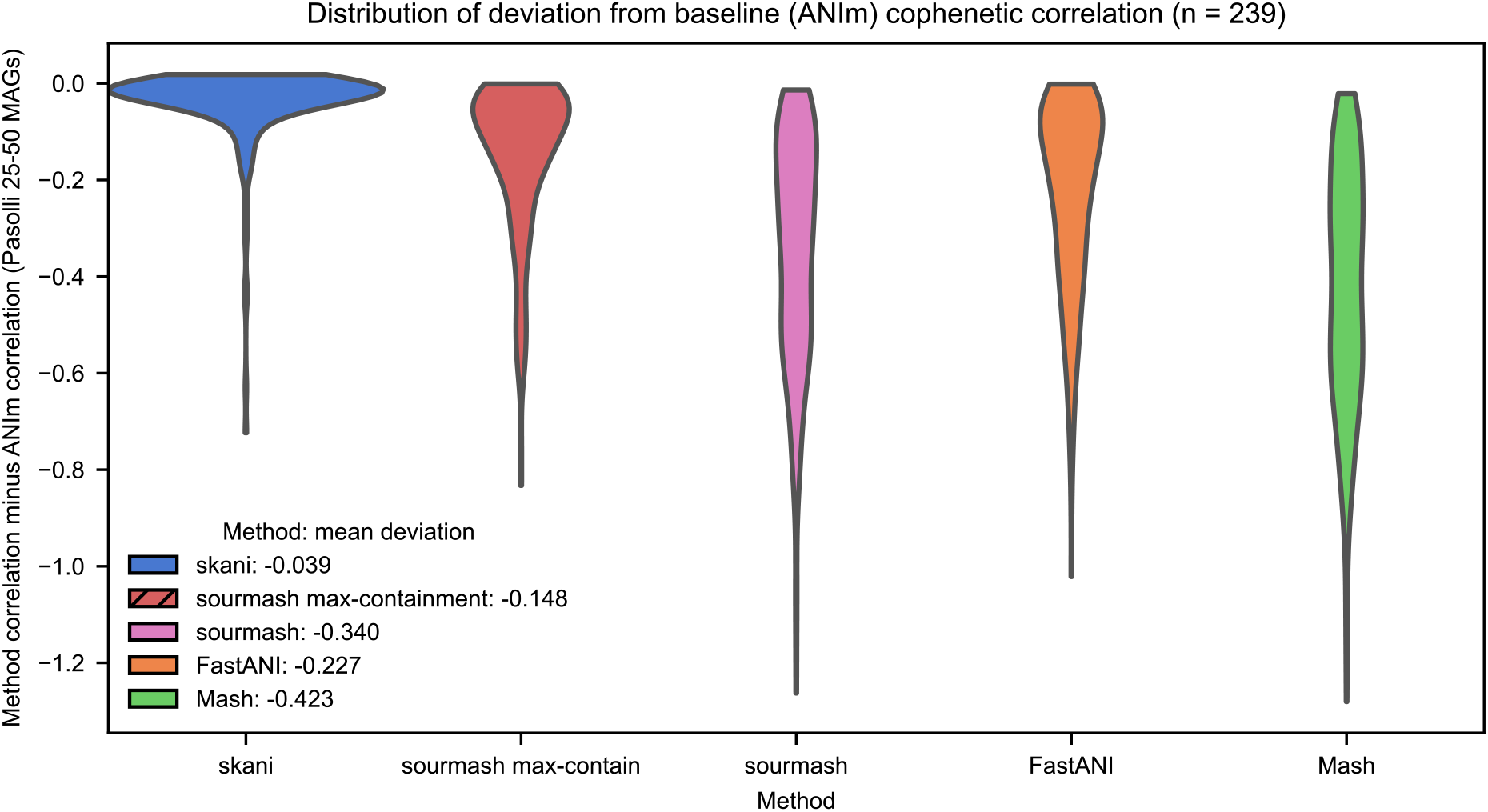
Violin plot of cophenetic correlation deviation for each method with respect to ANIm on the Pasolli et al 25-50 data set, where we took all species-level bins in Pasolli et al. [2] with > 25 and < 50 genomes and computed distance matrices for all ANI methods. sourmash max-contain implies the – max-containment option in sourmash, otherwise sourmash was run with default parameters. For each bin, we compared the resulting clustering dendrograms for each method against ANIm’s distance matrix by taking the cophenetic correlation. We plot the difference between this cophenetic correlation and the cophenetic correlation of ANIm’s dendrogram to itself (which may be < 1 since ANIm’s average-cluster dendrogram may not be perfectly concordant with itself). The legend shows the mean deviation of each method’s cophenetic correlation from ANIm’s cophenetic correlation.

**Extended Data Fig. 7:**
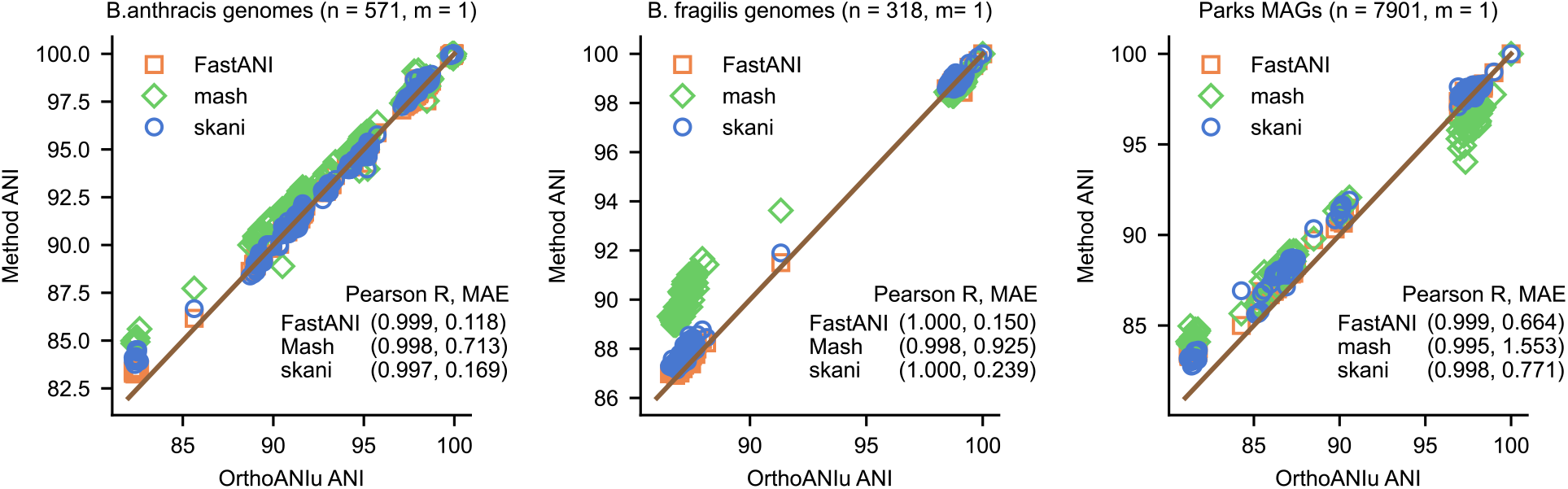
Additional ANI experiments on three different data sets specified in Supplementary Table 2. A single query genome (m = 1) was queried against n reference genomes. The same metrics and methodology was used as in Fig. 2 a.

**Extended Data Fig. 8:**
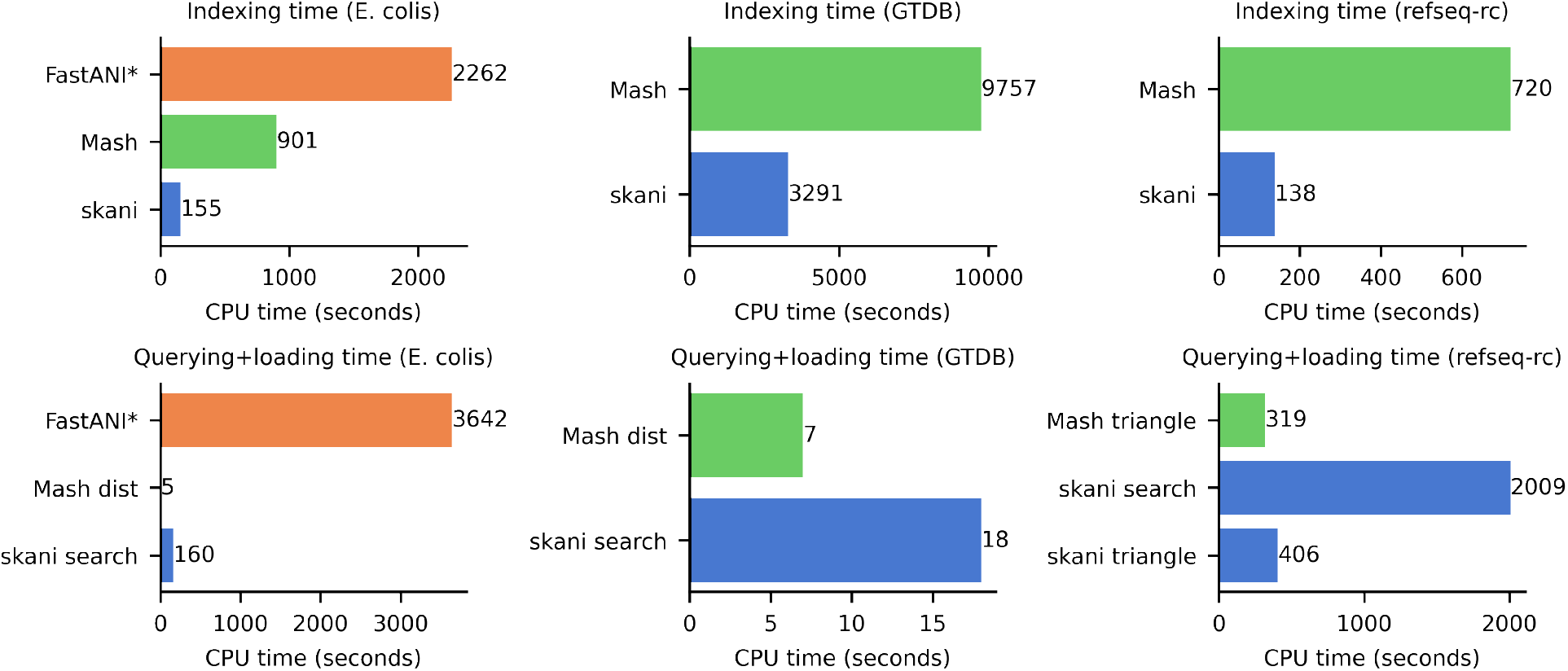
Benchmarking times for the three experiments run in Fig. 2 timed in CPU time. The exact same experiments were run, only with CPU times shown instead of wall times.

**Extended Data Fig. 9:**
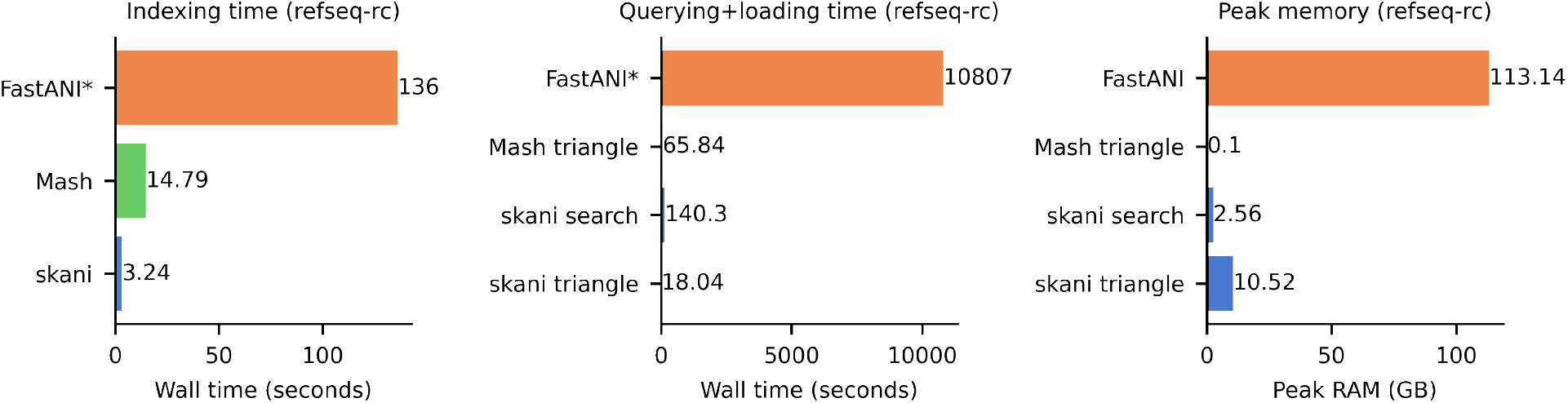
The same runtime as in Fig. 2 for the refseq-cr data set but with FastANI’s runtimes included. The timing comparison is unfair to FastANI because FastANI computes ANI for genomes as low as 75% ANI in practice and is more sensitive than skani, which uses an 80% putative ANI filter by default.

**Extended Data Fig. 10:**
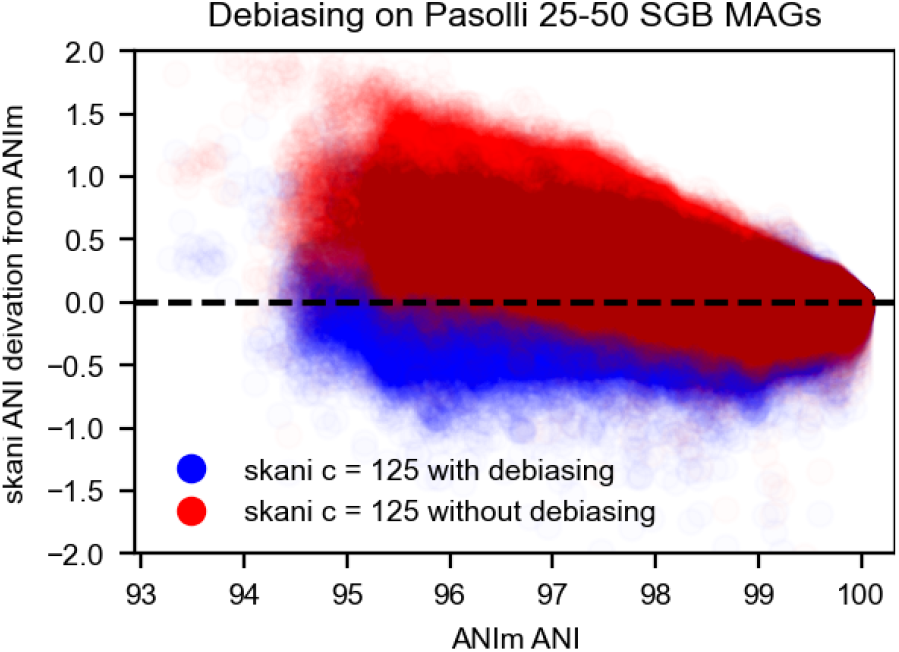
The effect of skani’s debiasing procedure for MAGs with the parameter *c* set to 125 (default). The y-axis shows the deviation from ANIm’s ANI estimate. This data set consists of all pairwise MAG ANIs within species level bins from Pasolli et al with between 25 and 50 genomes. Note the regression model is *not trained on these MAGs* but on a disjoint set of MAGs.

**Extended Data Fig. 11:**
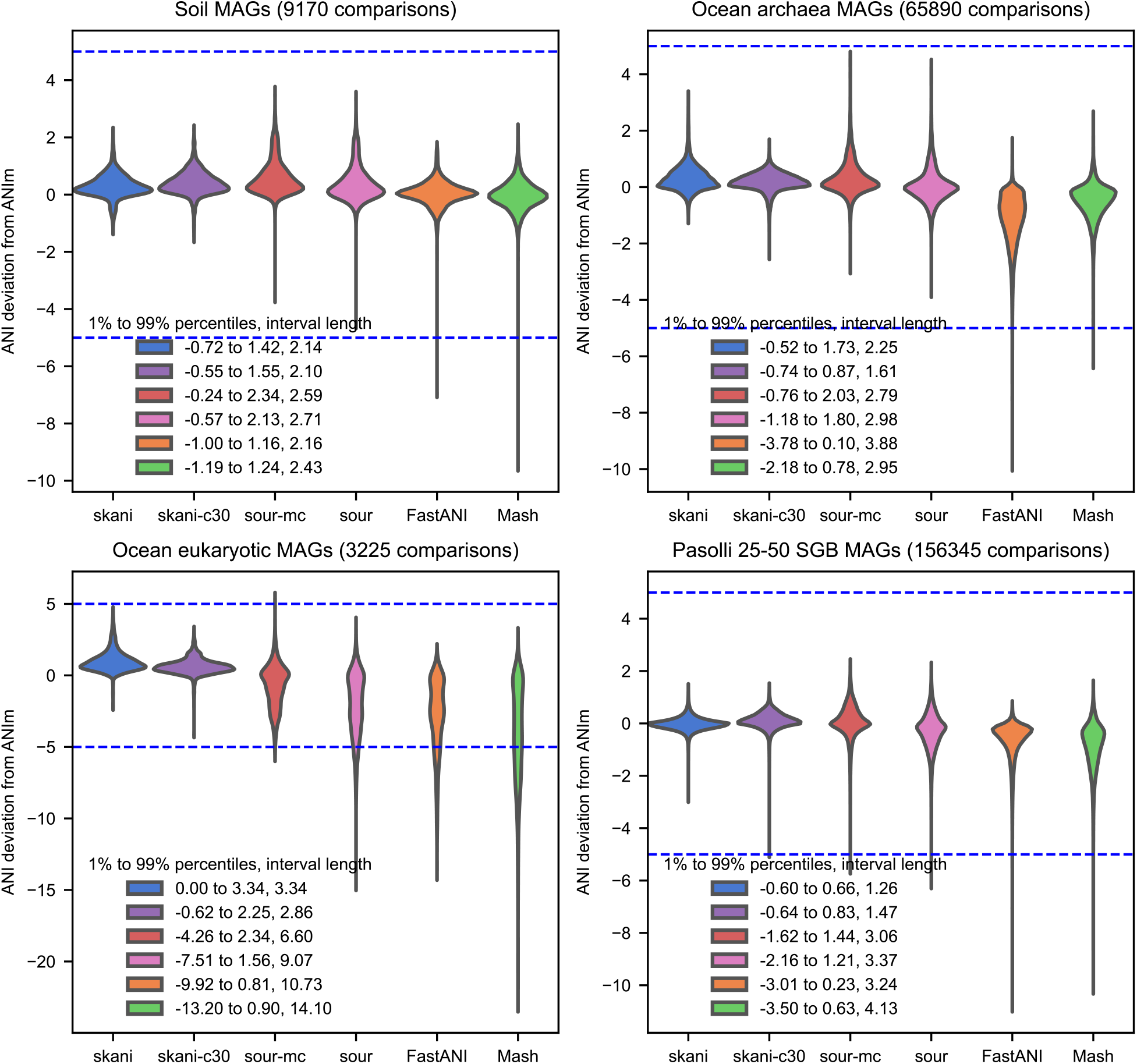
ANI deviation from ANIm for four data sets of MAGs with ¿ 90% ANI predicted by ANIm. See Supplementary Table 2 for data set descriptions. skani c-30 indicates skani with the parameter *c* set to 30. sour-mc and sour correspond to sourmash with max-contain enabled and disabled. Blue lines show 5 and -5 deviation, and the legend indicates the 1 and 99 percentile deviations as well as their difference. The best 1% to 99% deviation distance for each data set is either skani or skani-c30. Lowering the *c* parameter generally improves results slightly, but is not guaranteed to do so; the default *c* = 125 gives results more in line with ANIm on the Pasolli 25-50 data set.

**Extended Data Fig. 12:**
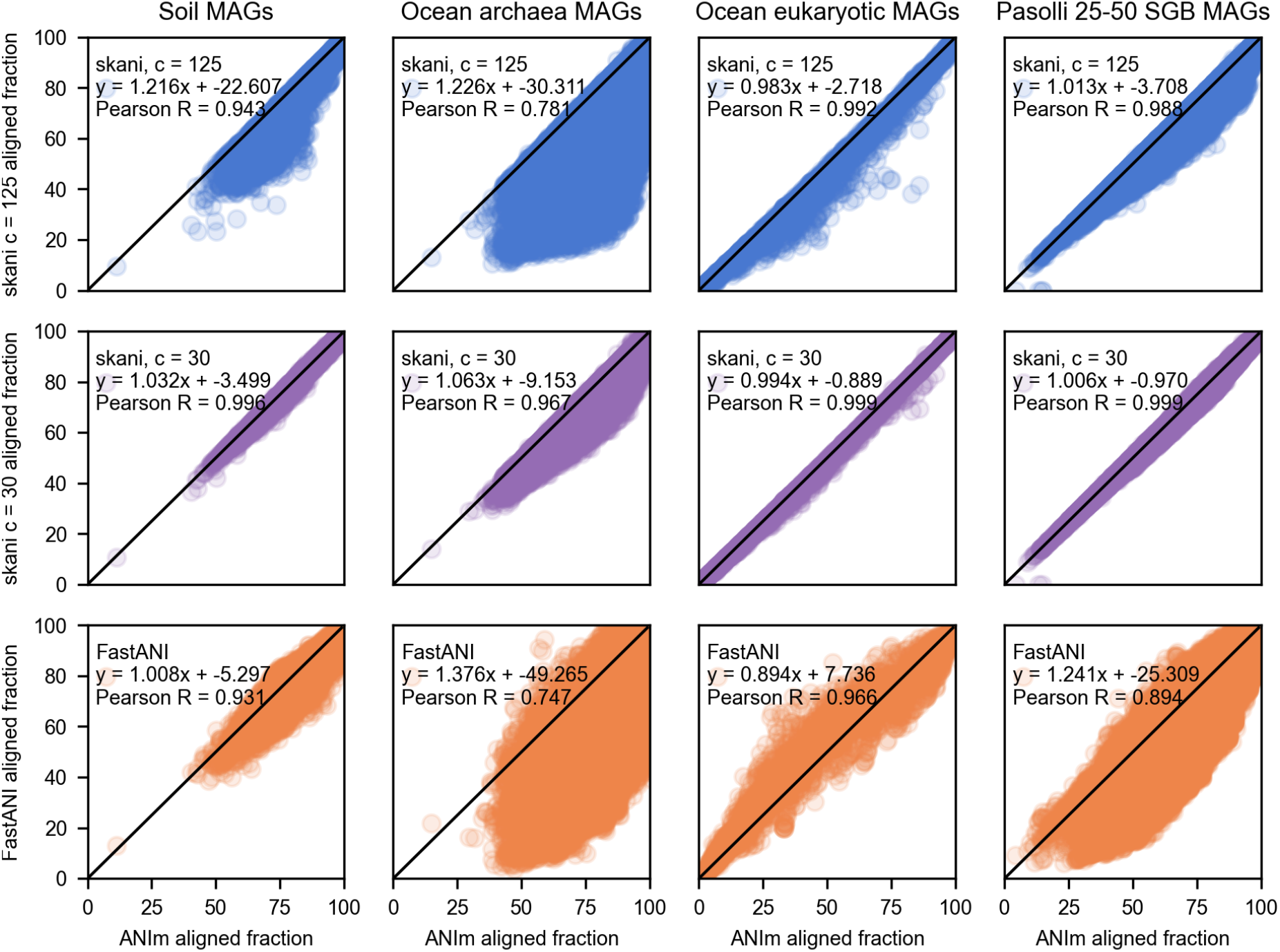
Aligned fraction correlation between ANIm and skani with default parameters (c = 125), skani with c = 30, and FastANI. Only MAGs with > 90% ANI as predicted by ANIm are compared. Lowering the value of *c* for skani gives a more accurate signal. skani with default parameters still outperforms FastANI in terms of Pearson R for all data sets, and skani with c = 30 has near perfect correspondence with ANIm. The noisier results for archaea MAGs are likely due to its small N50 (median 5863 bp) and genome length (median 1.27 Mbp).

**Extended Data Fig. 13:**
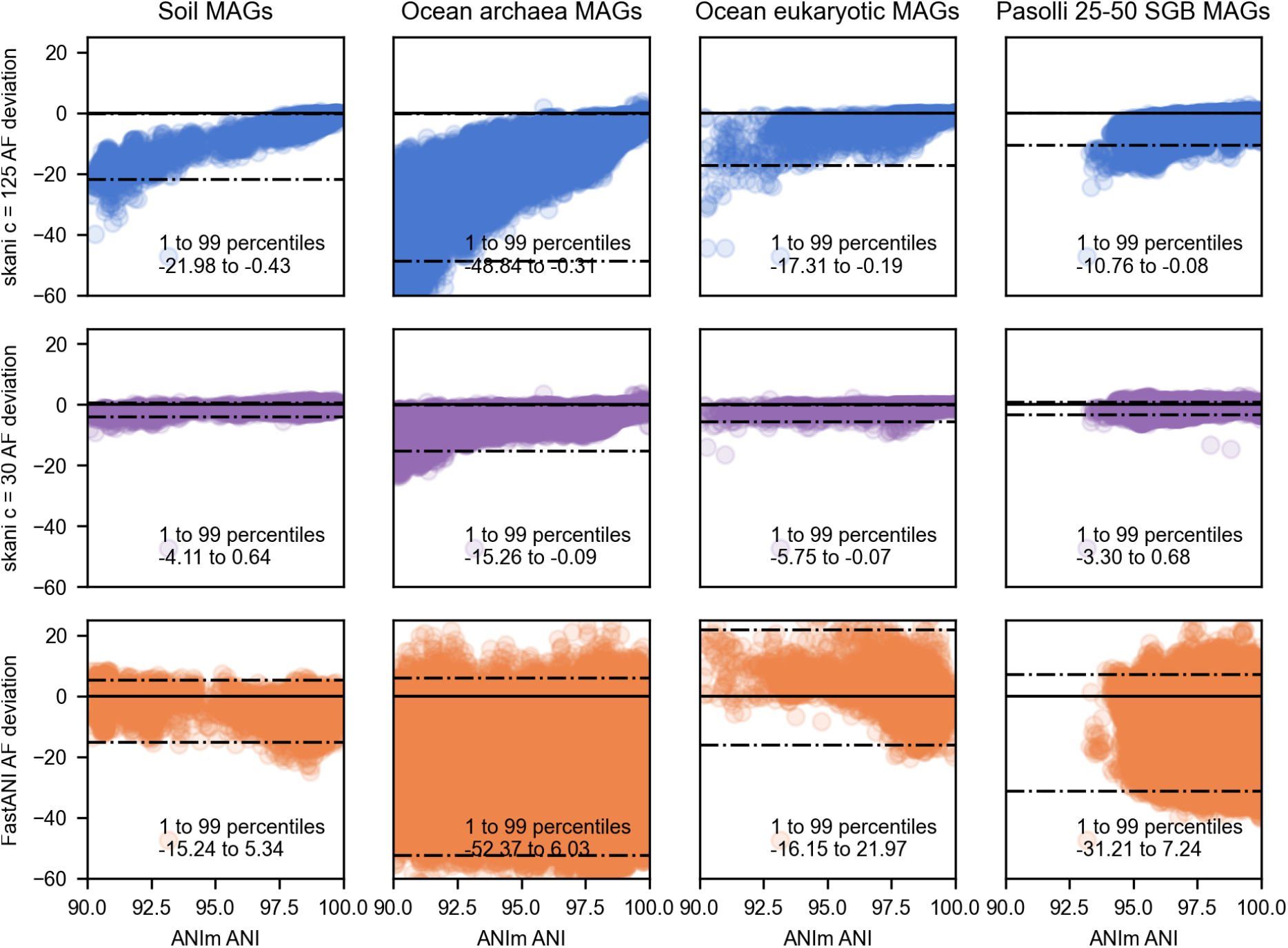
Aligned fraction deviation from ANIm as a function of ANI. Dashed lines indicate the 1 and 99 percentiles of aligned fraction deviation from ANIm. Aligned fraction gets more accurate for skani as ANI increases. Decreasing *c* for skani improves the relationship between ANI and aligned fraction and decreases the variance as well.

**Extended Data Fig. 14:**
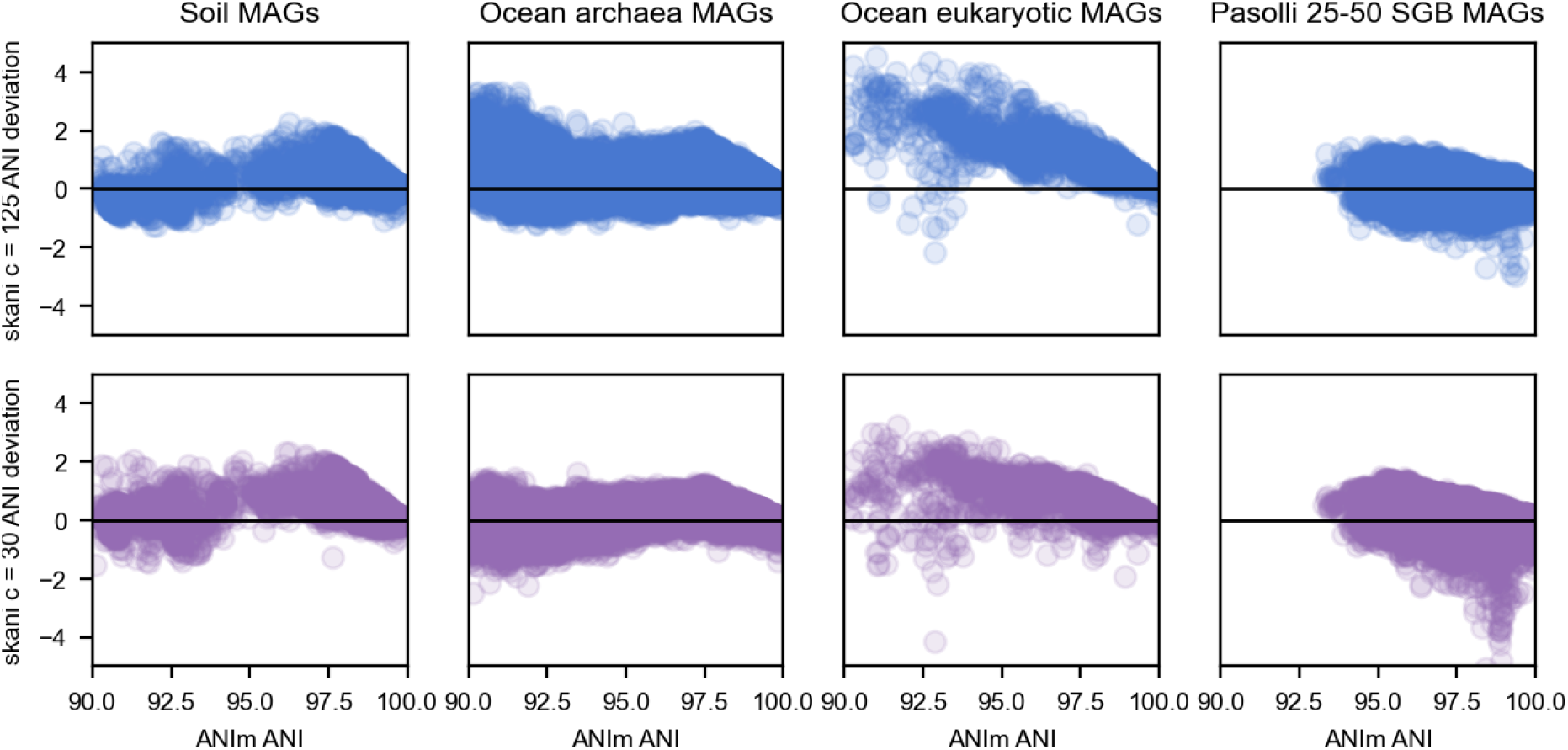
skani’s ANI deviation from ANIm as a function of ANI. On the eukaryotic and archaea data sets, lowering *c* visibly decreases the bias, especially for smaller ANI values close to 90%. However, it has a smaller effect on the soil and Pasolli data sets.

